# The *B. subtilis* Rok protein compacts and organizes DNA by bridging

**DOI:** 10.1101/769117

**Authors:** L. Qin, A.M. Erkelens, D. Markus, R.T. Dame

**Affiliations:** Leiden Institute of Chemistry, Leiden University, Einsteinweg 55, 2333CC Leiden, The Netherlands; Centre for Microbial Cell Biology, Leiden University, Leiden, The Netherlands

**Keywords:** bacterial chromatin, H-NS family proteins, DNA bridging, environmental sensing, adaptation

## Abstract

Rok from *Bacillus subtilis* is an abundant DNA binding protein similar in function to H-NS-like proteins found in many proteobacteria. Rok binds across the genome with a preference for A/T rich DNA. Such DNA often contains genes of foreign origin that are silenced due to Rok binding. Rok also has been implied in global organization of the *B. subtilis* genome. However, how Rok binds to DNA and how it represses transcription is unclear. Also, it is unknown whether Rok-mediated gene repression can be induced or relieved following changes in physico-chemical conditions, as noted for H-NS-like proteins. Here we investigate the DNA binding properties of Rok and determine the effects of physico-chemical conditions on these properties. We demonstrate that Rok is a DNA bridging protein similar to H-NS like proteins from *E. coli* (H-NS), *Pseudomonas sp.* (MvaT) and *Mycobacteria* (Lsr2). Strikingly, unlike these proteins, the ability of Rok to bridge DNA is not affected by changes in physico-chemical conditions. Not being a direct sensor of such changes sets Rok apart from other H-NS like proteins. It implies the existence of other (protein-mediated) mechanisms to relieve Rok-mediated gene silencing in response to changes in environmental conditions.

## Introduction

The bacterial genome, like that of eukaryotic cells, is both functionally organized and compactly folded. Nevertheless, genes need to be accessible for the transcription machinery or need to be made accessible if environmental conditions so require: the nucleoid is dynamically organized and re-organized [1, 2]. Many factors contribute to the compact shape of the nucleoid, including DNA supercoiling, macromolecular crowding and nucleoid-associated proteins (NAPs) [3-5]. The Histone-like nucleoid structuring protein (H-NS), one of the main NAPs in *Escherichia coli*, plays important roles in both genome organization and gene regulation [6]. H-NS non-specifically binds DNA across the genome, but has a preference for AT-rich DNA. DNA acquired via horizontal gene transfer (HGT) is often AT-rich and is recognized as xenogeneic DNA by H-NS [7, 8]. Although genes acquired via HGT are key to the evolution of bacteria by conferring new genetic traits, inappropriate expression of newly acquired genes can lead to loss of competitive fitness of bacteria. H-NS family proteins including H-NS of *E. coli*, MvaT of *Pseudomonas sp.* and Lsr2 of *Mycobacteria*, function as xenogeneic silencers, silencing foreign DNA until an environmental signal leads to relief of repression.

Akin to H-NS, MvaT and Lsr2 are also regulators of global gene expression. Although by sequence the proteins are not homologous, they are similar in DNA binding properties and function, due to their similar organization in functional domains. Structural studies have revealed that H-NS, Lsr2 and MvaT have an N-terminal oligomerization domain consisting of two dimerization sites, a C-terminal DNA binding domain and a flexible linker region [7, 9-12]. Due to their ability to dimerize and oligomerize, these proteins can bind along DNA forming a nucleoprotein filament, which stiffens DNA [13-15]. Under appropriate physico-chemical conditions the proteins also can bridge remote segments along a DNA duplex [16-18], yielding DNA loops. The bridging activity of H-NS and MvaT can be modulated by both monovalent (Na^+^, K^+^) and divalent (Mg^2+^, Ca^2+^) salt [19-22], while for Lsr2 it remains unclear whether ionic conditions affect its binding properties. The DNA binding activity of H-NS family proteins (either the formation of nucleoprotein filaments or bridged protein-DNA complexes) is sensitive to temperature, pH and salt, which contributes to adaptation of cells to environmental challenges, mediated by the bacterial genome [2, 6, 23]. Both filament formation and DNA bridging activity of H-NS family proteins have been suggested to account for gene regulation [20]. However, as only the DNA bridging activity can be switched on or off by small changes in physicochemical condition, this might be the mode of binding essential to regulation of environmentally regulated genes [24].

Rok of *Bacillus subtilis* was recently proposed to be a functional homolog of H-NS based on the observation that Rok binds extended regions along the genome, which are preferentially AT-rich and have been acquired via horizontal gene transfer. Rok contributes to silencing of the genes within such regions [25]. The ability of Rok to silence genes on DNA of foreign origin classifies it as a xenogeneic silencer like H-NS, MvaT and Lsr2. Rok is also found associated with a large subset of chromosomal domain boundaries in *B. subtilis* [26], which suggests it contributes to genome organization of *B. subtilis*. However, how Rok binds to DNA and how it silences genes remains unknown. In addition, it is unknown whether gene silencing by Rok can be modulated by changes in physico-chemical growth conditions such as temperature, pH and salt.

### Rok compacts DNA

All H-NS family proteins (H-NS, MvaT and Lsr2) exhibit two modes of binding to DNA: filament formation along DNA and DNA bridging [14, 15, 17-19, 22]. Both lateral nucleoprotein filament complex formation and DNA bridging are suggested to be important for the function of these proteins in gene silencing. To probe whether Rok, like H-NS family proteins, exhibits DNA stiffening activity (reflecting the formation of a protein filament along DNA) or not, we investigated the effect of Rok binding on the conformation of DNA by using Tethered Particle Motion (TPM) [27, 28]. Here, the Root Mean Square displacement (RMS) of a bead (exhibiting thermal motion) at the extremity of a DNA substrate attached to a glass surface, gives a readout of DNA conformation. If a protein stiffens DNA, the RMS will increase following the binding of proteins. If a protein softens, bends, or bridges DNA, a reduction in RMS will take place. We investigated the interaction between Rok and an AT-rich (32%GC) DNA substrate, which we used earlier in studies of the DNA-binding properties of H-NS [21] and MvaT [22]. We determined the effect of Rok on DNA conformation by titration from 0-10 nM (Figure 1a). Bare DNA has an RMS of 150 ± 2 nm. Upon addition of increasing amounts of Rok, a second unique population at an RMS of about 80 nm appears. Saturation of Rok binding is achieved at 8 nM; at this concentration only the population with reduced RMS is observed (Figure 1b). The observed reduction of RMS is an indication of DNA compaction by binding of Rok; it implies that Rok does not form filaments along DNA as observed for other H-NS-family proteins under similar conditions [14, 15, 19] (Figure 1c). The fact that compaction occurs at low protein concentration and that the structural transition is abrupt is suggestive of cooperative behavior. The reduction in RMS can be attributed either to DNA bending as observed for HU (Figure 1c) or to DNA bridging.

**Figure 1.**
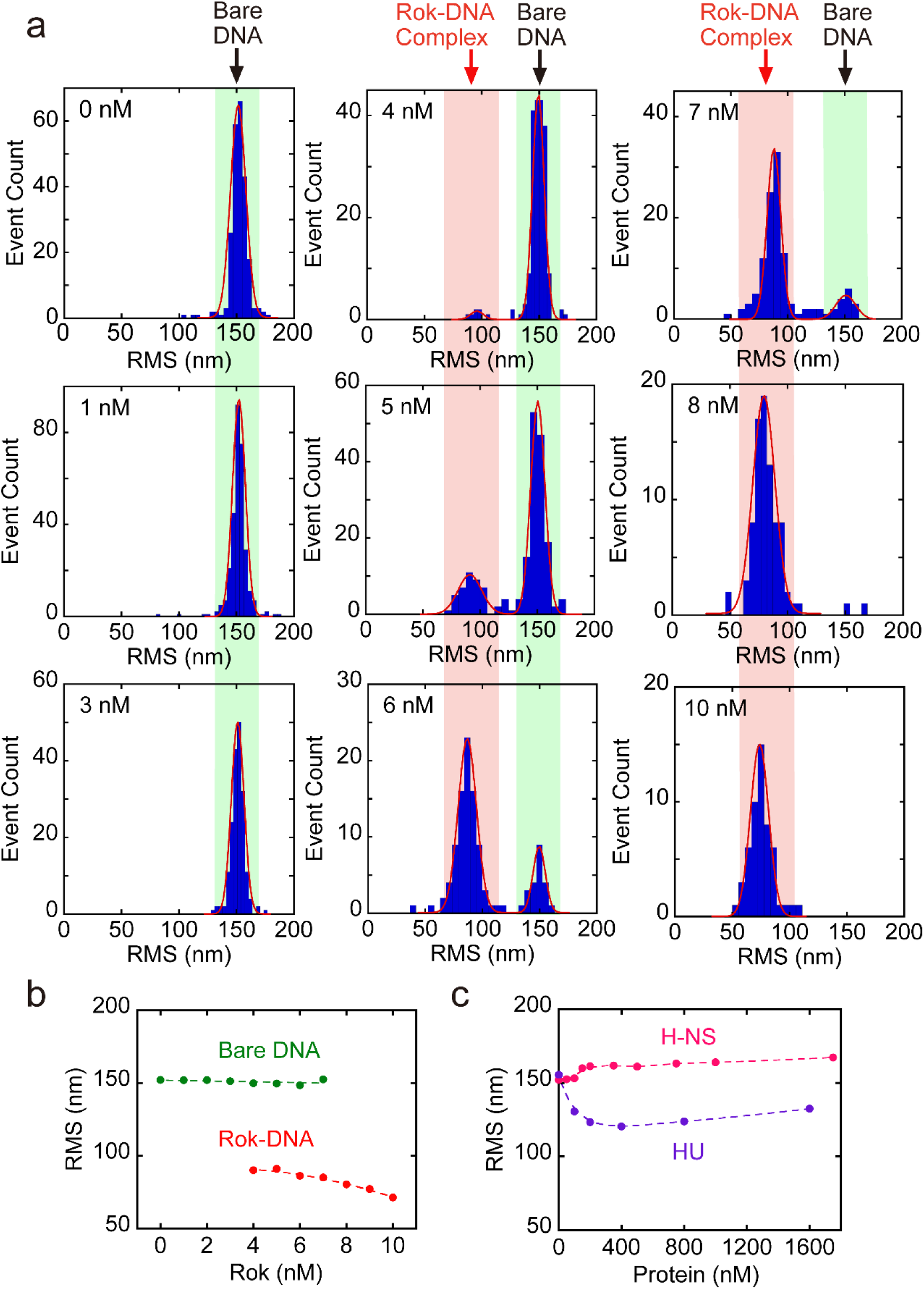
*B. subtilis* Rok compacts DNA. a) Histograms of Root Mean Square (RMS) obtained for 32%GC DNA as a function of Rok at concentrations of 0, 1, 3, 4, 5, 6, 7, 8, and 10 nM as measured by TPM in the presence of 50 mM KCl. The histograms were fitted to Gaussian distributions, in which the RMS value at around 150 nm represents bare DNA and the population of RMS at around 80 nM represents DNA bound by Rok. The bare DNA and Rok-DNA complex populations are highlighted with a light green and red box, respectively. The data for each concentration is the combination of three independent measurements and the RMS for each concentration was obtained by fitting the combined data to a Gaussian distribution. b) RMS values obtained for 32%GC DNA as a function of Rok at concentrations from 0 nM to 10 nM. Green and red dots represent the average RMS resulted from fitting with a Gaussian distribution, where green dots and red dots represent bare DNA and Rok-DNA complexes, respectively. Error bars represent the standard error of the mean; due to their small size they are hidden behind the data points. How the RMS values are distributed among the two populations is not taken into account in this representation. c) RMS as a function of protein concentration of *E. coli* H-NS and HU. Error bars represent the standard error of the mean; due to their small size they are hidden behind the data points. The Rok coding gene from *B. subtilis* was cloned into pET30b using Gibson Assembly [29] resulting in vector pRD231. Following overproduction in Rosetta™ (DE3) pLysS, cells were lysed and the lysate was centrifuged for 30 min at 37000 rpm. The supernatant was filtered with a 0.22 µm Millex-GP Syringe Filter. Next, the protein was purified using a HiTrap Heparin HP 1 mL affinity column (GE Healthcare), a HiTrap SP HP 1 mL column (GE Healthcare) and a GE Superdex 75 10/300 GL column. The purified protein was checked by mass spectrometry. The concentration was determined using a Pierce™ BCA Protein Assay Kit (Thermo Scientific). The DNA used for Tethered Particle Motion (TPM) experiments is a random, AT-rich (32% GC), 685 bp DNA substrate [21, 30]. Measurements were performed as previously described [21] with minor modifications. Briefly, the flow cell was washed with 100 µL experimental buffer (10 mM Tris-HCl pH 8, 10 mM EDTA, 5% glycerol, 50 mM KCl) to remove excess beads and 100 µL protein diluted in experimental buffer is flowed in and incubated for 5 minutes. Next, the flow cell was washed with protein solution one more time, sealed with nail polish and incubated for 5 minutes. After incubation, the flow cell was directly transferred to the holder and incubated for 5 more minutes to stabilize the temperature at 25°C for the measurement. For each flow cell more than 200 beads were measured and measurements for each concentration were performed in triplicate.

### Rok is able to bridge DNA

In order to determine the structural basis of the observed DNA compaction, we next investigated the ability of Rok to bridge DNA in a quantitative biochemical DNA bridging assay, which we used earlier to evaluate the impact of buffer conditions on the DNA bridging efficiency of H-NS and MvaT [21, 22]. In this assay biotinylated DNA is bound to streptavidin-coated beads and radioactively labeled DNA offered in trans can be recovered by magnetic pull-down of beads when bridged by protein. The radioactive signal of the DNA pulled down is a proxy of DNA bridging efficiency. To determine whether Rok bridges DNA we carried out a titration with Rok from 0 - 0.5 μM. The DNA (685 bp, 32% GC) used in the bridging assay is same as the DNA used in TPM experiments. In the absence of Rok, no radioactive DNA was recovered. DNA recovery increases upon addition of increasing amounts of Rok. Saturation of DNA recovery occurs at a Rok concentration of 0.3 μM (figure 2). These data unambiguously show that Rok is a DNA bridging protein. Rok has a similar DNA bridging efficiency as H-NS, yet reaches this efficiency at 10 times lower concentration (0.3 μM vs 3 μM) [21], which is attributed to the high cooperativity in DNA binding by Rok.

**Figure 2.**
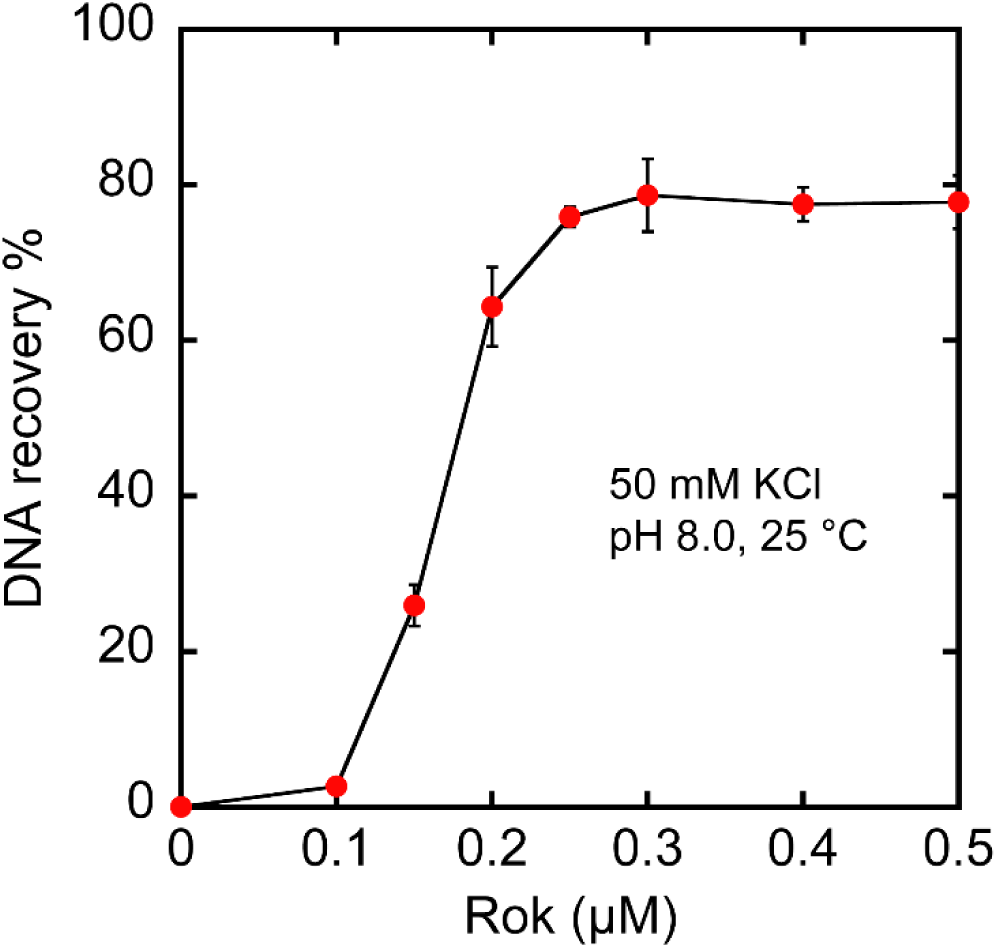
Rok bridges DNA. A) DNA recovery (as a percentage of the input DNA) as a function of Rok concentration from 0 to 0.5 µM as measured by the DNA bridging assay in the presence of 50 mM KCl at 25 °C. Data are plotted as mean values and the error bars represent the standard deviation from independent triplicate measurements. The DNA used for the bridging assay, the same as that used for TPM, was ^32^P-labeled as described previously [31]. The DNA bridging assay was performed as previously described [21, 32] with minor modifications. Streptavidin-coated Magnetic M-280 Dynabeads (Thermo Fisher) were resuspended in buffer (20 mM Tris-HCl pH 8.0, 2 mM EDTA, 2 M NaCl, 2 mg/mL BSA (ac), 0.04% Tween 20) containing 100 fmol biotinylated 32% GC DNA (685 bp) and incubated at 1000 rpm for 20 min at 25°C. The beads with associated DNA were washed twice before resuspension in buffer (10 mM Tris-HCl, pH 8.0, 5% v/v Glycerol, 1 mM Spermidine). Radioactive ^32^P-labeled DNA and unlabeled DNA were combined to maintain a constant (2 fmol/μL) concentration and a radioactive signal around 8000 cpm, and then added to each sample. Next, Rok was added (concentration as indicated) to initiate formation of bridged protein-DNA complexes. The samples were incubated for 20 min at 1000 rpm at 25°C. After the incubation, the beads were washed with the same experimental buffers before resuspension in counting buffer (10 mM Tris-HCl, pH 8.0, 1 mM EDTA, 200 mM NaCl, 0.2% SDS). The radioactive signal of DNA was quantified by liquid scintillation and was used for the calculation of protein DNA bridging efficiency (%) based on a reference sample containing the same amount of labeled ^32^P 685 bp DNA used in each sample. All DNA bridging experiments were performed at least in triplicate.

### DNA bridging activity of Rok is not sensitive to environmental conditions (temperature, pH and salt)

Bacteria adapt to environmental changes and environmental cues have a direct effect on the function of H-NS-like proteins. This is suggestive of environment-sensory activity of these proteins at H-NS-regulated environment-responsive genes. The DNA bridging activity of H-NS is sensitive to environmental conditions, such as salt [19]. Adaptation of *B. subtilis* to osmotic up- and downshift is a frequent challenge, which is also essential for growth and survival in its natural living environment [33]. To determine whether Rok’s DNA bridging activity is also sensitive to salt condition, we investigated the effect of changing monovalent and divalent salt concentration. An increase in concentration of KCl from 50 mM to 300 mM has no significant effect on the DNA bridging activity of Rok. Nevertheless, at higher concentration of KCl the formation of bridged complexes is abolished (figure 3a). A similar result was obtained for titration with MgCl_2_ in the range from 0 mM to 90 mM (figure 3b). Only beyond a certain ionic strength, the bridged Rok-DNA complexes start disintegrating and this ionic strength is likely not physiologically relevant anymore. It has been reported that the bridging activity of H-NS and MvaT can be modulated by both monovalent (K^+^) and divalent (Mg^2+^) salt [19-22]. However, unlike H-NS and MvaT, the formation of bridged Rok-DNA complexes requires neither Mg^2+^ nor a high concentration of K^+^. These results indicate that DNA bridging activity of Rok neither requires nor is inhibited by salt at biologically relevant concentrations.

**Figure 3.**
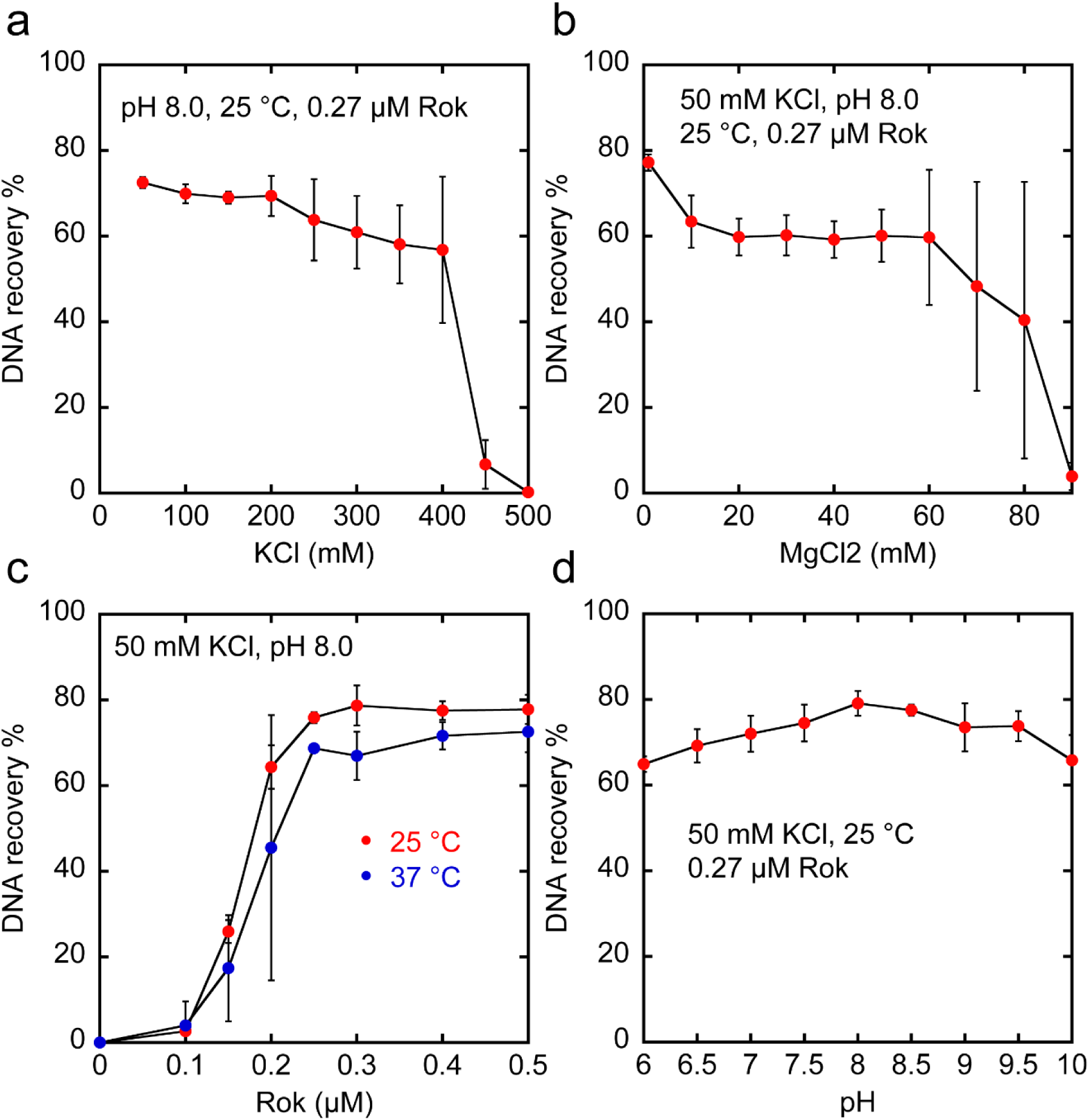
DNA bridging activity of Rok is not sensitive to temperature, pH and salt. A) DNA recovery (as a percentage of the input DNA) as a function of KCl concentration from 50 to 500 mM measured by the DNA bridging assay in the presence of 0.27 µM Rok at 25 °C. b) DNA recovery as a function of MgCl_2_ concentration from 1 to 90 mM in the presence of 0.27 µM Rok at 25 °C. c) DNA recovery as a function of Rok concentration from 0 to 0.5 µM at 25 °C (red) and 37 °C (blue), respectively. d) DNA recovery as a function of pH from 6 to 10 in the presence of 0.27 µM Rok at 25 °C. Data are plotted as mean values and the error bars represent the standard deviation from three independent measurements. The experiment was carried out in the same way as described for figure 2. Salt concentration (KCl or MgCl_2_), protein concentration, temperature and pH were varied in line with the experiments. For pH 6 and 6.5 MES (2-morpholinoethanesulfonic acid) was used instead of Tris-HCl and for pH 9, 9.5 and 10 CHES (*N*-Cyclohexyl-2-aminoethanesulfonic acid) was used.

*B. subtilis*, found in soil and the gastrointestinal tract of ruminants and humans, can live at different temperatures and pH conditions. Therefore, we investigated the effect of temperature on DNA bridging efficiency. An increase in temperature from 25 °C to 37 °C has no significant effect on the DNA bridging activity of Rok (figure 3c). Next, we also investigated the effect of pH. Across a range from 6 to 10, pH has no effect on the DNA bridging activity of Rok (figure 3d). Strikingly, even crossing the pI of Rok (9.31) did not interfere with its capacity of DNA bridging. Taken together, all these results indicate that the DNA bridging activity of Rok is not affected by changes in physico-chemical conditions, which is unexpectedly different from H-NS, MvaT and Lsr2.

We have studied the DNA binding properties of Rok. We found that Rok is a DNA bridging protein and evaluated its DNA bridging capacity under various physiologically relevant conditions. The protein is remarkably insensitive to changes in physico-chemical conditions. Although Rok has low sequence similarity with H-NS, MvaT or Lsr2, they have similar domain organization: the N-terminal dimerization and oligomerization domain, a C-terminal DNA binding domain and a flexible linker in between. H-NS, MvaT and Lsr2 can oligomerize along DNA, which causes DNA stiffening. Such a stiffening effect was not observed for Rok. Instead, only DNA compaction was observed in our study. Because we demonstrated that Rok is a DNA bridging protein, we attribute the observed DNA compaction to bridging by Rok. Based on our results and the known properties of Rok, we propose that Rok bridges DNA, without associating into nucleoprotein filaments, but employing dimeric Rok as bridging units (Figure 4). The Rok dimers cluster cooperatively due to high local DNA concentration upon initiation of bridging, but earlier studies suggest that the protein forms oligomers in solution [34], while our study does not provide evidence for Rok oligomers along DNA. Unexpectedly, Rok-DNA bridging activity is not affected by changes in physico-chemical conditions, which sets the protein apart from H-NS, MvaT and Lsr2. It implies the presence of other (protein-mediated) mechanisms to relieve Rok-mediated gene silencing in response to changes in environmental conditions. An example of a Rok antagonist is ComK which can relieve gene repression mediated by Rok at the *comK* promotor [35]. Based on the robustness of Rok in binding to DNA we expect the existence of similar antagonistic factors operating at other sites across the genome.

**Figure 4.**
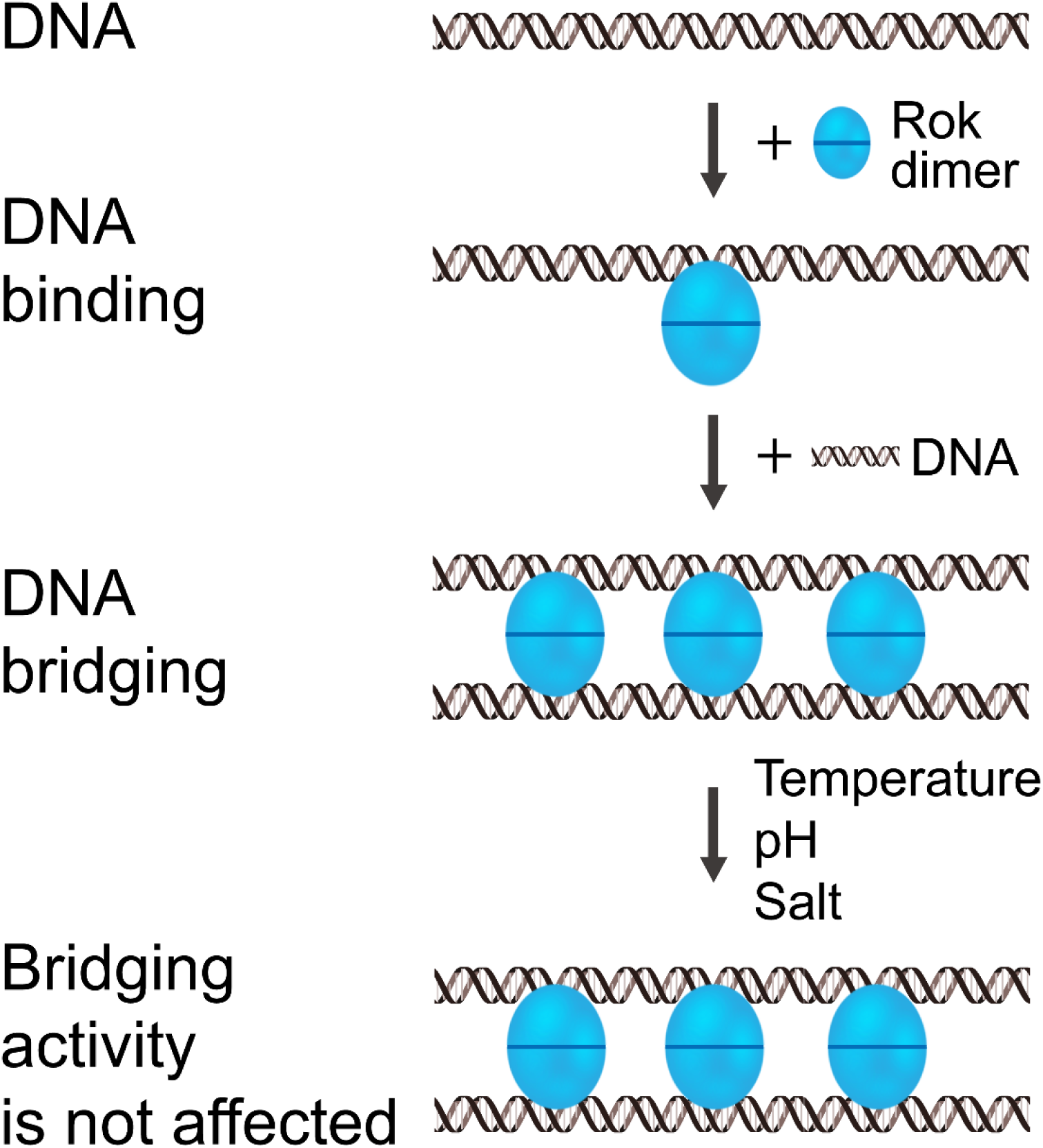
The proposed mechanisms of DNA binding by Rok. Rok binds DNA as a dimer without associating into nucleoprotein filaments. Dimeric Rok acts as bridging unit. Rok dimers cluster cooperatively in between two DNA duplexes, not due to dimer-dimer interactions, but due to high local DNA concentration which drives association and bridging by additional dimers. The DNA bridging activity of Rok is not sensitive to changes in physico-chemical conditions (temperature, pH and salt).

## Acknowledgments

This research was supported by grants from the Netherlands Organization for Scientific Research [VICI 016.160.613] (R.T.D.), the Human Frontier Science Program (HFSP) [RGP0014/2014] (R.T.D.) and a grant from the China Scholarship Council (CSC) [201506880001] (L.Q.).

## Contributions

L.Q., A.M.E. and D.M. performed the experiments. L.Q., A.M.E., D.M. and R.T.D. contributed to data analysis and discussion. L.Q. and R.T.D. supervised the project. L.Q., A.M.E., and R.T.D. wrote the manuscript. L.Q., A.M.E. and R.T.D. reviewed and corrected the manuscript.

